# Agarose disk electroporation method for *ex vivo* retinal tissue cultured at the air-liquid interface reveals electrical stimulus-induced cell cycle reentry in retinal cells

**DOI:** 10.1101/2023.12.21.572865

**Authors:** Megan L. Stone, Hannah H. Lee, Edward M. Levine

**Affiliations:** Department of Cell and Developmental Biology Vanderbilt University, Nashville TN 37232; Department of Ophthalmology and Visual Sciences Vanderbilt University Medical Center, Nashville TN 37232

## Abstract

It is advantageous to culture the *ex vivo* murine retina along with many other tissue types at the air-liquid interface. However, gene delivery to these cultures can be challenging. Electroporation is a fast and robust method of gene delivery, but typically requires submergence in a liquid buffer to allow electric current flow. We have developed a submergence-free electroporation technique using that allows for efficient gene delivery to the *ex vivo* murine retina. This method advances our ability to use *ex vivo* retinal tissue for genetic studies and can easily be adapted for any tissue cultured at an air-liquid interface. Use of this method has revealed valuable insights on the state of *ex vivo* retinal tissues and the effects of electrical stimulation on retinal cells.

**Motivation:** Tissues cultured at the air-liquid interface, such as retinal tissue, are adhered to a filter membrane with media underneath but not fully submerged. If tissues are fully submerged in liquid, they detach from the membrane and become damaged. Electroporation typically requires tissue submergence for electric current flow. We have developed a submergence-free electroporation method using an agarose disk for electric current delivery that can efficiently and consistently electroporate *ex vivo* retinal tissue cultured at an air-liquid interface without the need to submerge the tissue and disrupt the culture system.

## Introduction

The retina is a highly structured organ that relies on intercellular communication to respond to injury and maintain homeostasis^1–3^. The field of retinal research relies on various model systems to study the structure, function, and pathology of the retina. *Ex vivo* explant systems involve isolating and culturing tissue samples, which allows for the preservation of the complex structure and physiological characteristics of organs like the retina^4–6^. *Ex vivo* retinal tissue cultures offer several advantages over other models, particularly in the study of retinal function and degeneration. *Ex vivo* models allow the study of cellular changes within the context of the tissue structure compared to *in vitro,* which lack the 3-dimensional context, or organoid models, which lack prior organismal physiological integration and external environmental exposures. Furthermore, *ex vivo* models provide the opportunity for more advanced manipulations and analyses that may be difficult to perform and interpret *in vivo*, such as live imaging, small molecule delivery and cell reprogramming studies^7,8^.

*Ex vivo* retinal tissues experience cell death and activation of Müller glia over time in culture. This makes *ex vivo* retinal explant models particularly valuable as an injury model^6,9–12^. It is of interest to improve the health of these cultures to study the retina under normal physiological conditions, however. Innovations in culture technique and supplementation has been shown to promote the survival of *ex vivo* retinal explants. One such innovation is to culture tissue on PTFE or polycarbonate membrane inserts with media below the membrane filter. This technique maintains an air-liquid interface that is valuable for gas exchange in the tissue^9,13,14^. Lung, liver, skin and brain tissue are commonly cultured at the air-liquid interface and the methods described here could be adapted for use with these tissues^15–19^. Optimization studies to preserve the health of *ex vivo* retinas are ongoing, but as is, long term culture of *ex vivo* retinal tissue is a useful injury context to observe live changes in response to genetic manipulation.

Gene delivery to *ex vivo* retinal tissue can be challenging. Viral methods, such as AAV and lentivirus, are limited by size constraints for DNA cargo and have delayed expression time compared to non-viral methods^20,21^. Electroporation is a robust method for plasmid delivery that promotes quick expression^22–24^. However, electroporation requires that cells or tissues be submerged in buffer with an electrode to allow electric current flow and that direct contact between the tissue and electrodes is avoided. Submersion of tissue is feasible prior to culture, but submerging tissues at the air-liquid interface causes detachment from the support membrane, which restricts the ability to transfect after cultures are established.

To overcome the obstacles to delivery of genetic material to *ex vivo* retinal tissues, we developed an electroporation method that avoids tissue submergence in a large volume and prevents direct contact with the electrode by generating a confined liquid interface between the electrode and sample (Figure 1). With this adaptation, we characterized the efficiency of electroporation at 1- and 14-days *ex vivo* (DEV) in the air-liquid interface culture format. We also characterized the effects of electrical stimulation on cell proliferation in *ex vivo* retinal cultures to provide clarity to investigators that may use this method in future studies that the effects of electroporation are not inert in the *ex vivo* retinal system. Lastly, we demonstrate the limitations in assessing transfection efficiency when relying on a fluorescent reporter construct as a readout.

**Figure 1.**
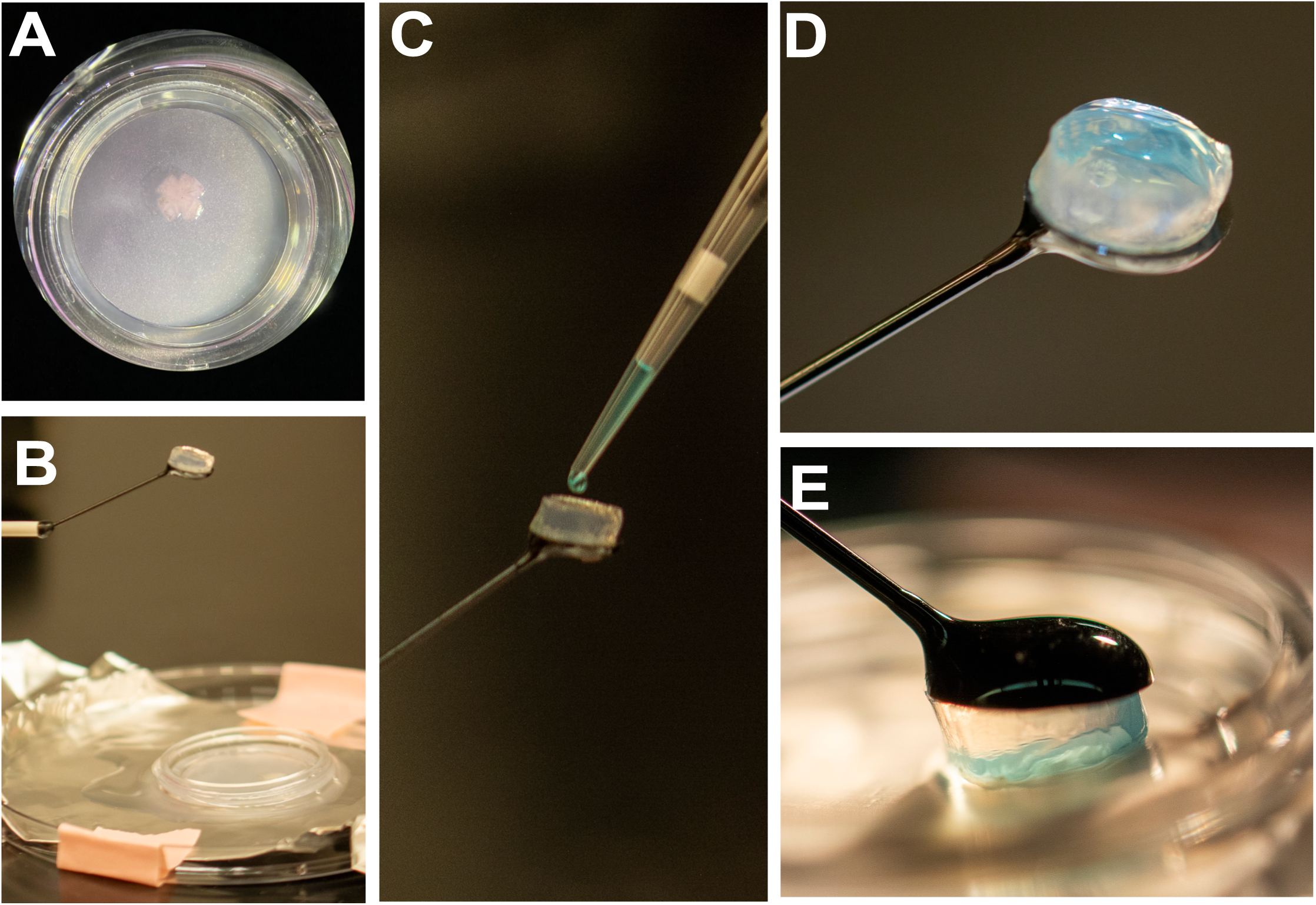
Visual methodology of agarose disk electroporation for *ex vivo* retinas. (A) Murine retinal tissue cultured at the air-liquid interface on PFTE membrane after 5 days *ex vivo* (5 DEV) (1X, brightfield). (B) Agarose disk electroporation set-up. 5mm high, 10mm diameter 0.5% agarose disk adhered to an arm electrode above an anode dish containing *ex vivo* retinal tissue on PTFE membrane with chilled PBS underneath. (C) 12uL of DNA-glycerol mixture is pipetted onto the agarose disk facing upward. (D) DNA-Glycerol mixture on top of agarose disk facing upward. (E) 180° rotation electrode arm containing agarose disk and DNA-glycerol mixture lowered onto *ex vivo* retinal tissue.

## Results

### Agarose disk electroporation of *ex vivo* retinal tissue efficiently delivers genetic material with minimal tissue submergence

The components required for this electroporation system are: (1) an arm electrode that can be fixed to a ring stand and lowered onto cultured retinas in cell culture insert, (2) a 0.5% agarose disk, 5mm thick and the width of the arm electrode, (3) DNA with an electroporation dye containing glycerol to increase viscosity, and (4) an anode dish; aluminum foil at the bottom of a 100mm petri dish, underneath the tissue. Electroporation was performed at 24 hours (1DEV), or 14 days (14DEV) and retinas were fixed at 72-96 hours post-electroporation. All *ex vivo* retinas were dissected from mice between 8-12 weeks of age.

Prior to the experiment, 500uL of 0.5% liquified agarose (RPI, A20090) is pipetted into 1 well of a 24-well plate (Corning, CLS3527) and set aside for 30 minutes in a sterile culture hood. A disk roughly 10 mm wide is created from the solidified agarose using the end of a plastic transfer pipet (Fisherbrand 13-711-7M), cut above the tapered tip, to stamp a disk shape. The agarose disk is then adhered to the arm electrode (Bulldog Bio, CUY700P7L) surface (Figure 1C) while the arm electrode is suspended face up using a micromanipulator (Narishige, UMM-3C). The disk should be adhered with no air bubbles formed between it and the electrode, which could impede electrical flow. Once the agarose disk is adhered to the electrode, 10-15uL of DNA-glycerol mixture (1-3 pMol DNA, 2.5% methyl green dye (MCE, HYD0163) and 7.5% glycerol (Sigma Aldrich, G7893) is pipetted on top of the agarose disk facing upward (Figure 1C, D). To complete the electric circuit, chilled PBS (Corning, 21-040-CV) (4°C) is placed on the anode surface and the cell culture insert (Millicell, PIC0RG50) containing the retinal tissue is placed on top (Figure 1A, B). The arm electrode with the agarose disk and DNA mixture is rotated 180 degrees and the DNA mixture will stay adhered to the agarose as a suspended drop. The arm electrode is lowered until the DNA mixture contacts the tissue, at which point current is delivered in five 50ms square wave pulses (25V) with 250ms intervals between pulses (Figure 1E). Fluorescent reporter-expressing plasmids with a general promoter, such as pCMV-eGFP, were used to assess transfection.

### Differences in cell types transfected at the edge and center of the tissue at 1DEV

Since retinas were cultured with their basal surface toward the air-liquid interface, the cells most accessible to plasmid uptake were astrocytes, retinal ganglion cells, displaced amacrine cells, and Müller glia^25,26^. Electroporation (referred to as transfection hereon) of pCMV-eGFP at 1DEV revealed a consistent difference in morphology from cells transfected at the center of the retina compared to the edge that is defined as 100µm in from tissue periphery around perimeter (Figure 2B). Cells transfected at the edge of the tissue resembled Müller glia in morphology (Figure 2D, G). Colocalization of eGFP with SOX9, a marker of Müller glia^27–29^, is consistent with this observation (Figure 2D). In contrast, transfected cells toward the center of the tissue had a retinal ganglion cell-like morphology, which is distinctly recognizable by long axons converging at the optic nerve head (Figure 2C, F). Transfection of pCMV-mCherry^30^ into retinas from Thy1:YFP-16 mice (Thy1:YFP), which express YFP in a large cohort of RGCs in addition to other inner retinal neurons^31^, provided further evidence that RGCs were transfected (Figure 2G). Quantifications revealed that the distributions of transfected cells were skewed toward SOX9+ cells at the tissue edge and toward Thy1:YFP+ cells toward the center (Figure 2H). Overall, there is equivalent efficiency in transfection of Müller glia and Thy1:YFP+ cells in retinas transfected at 1DEV, but their distribution is location-dependent (Figure 2H).

**Figure 2.**
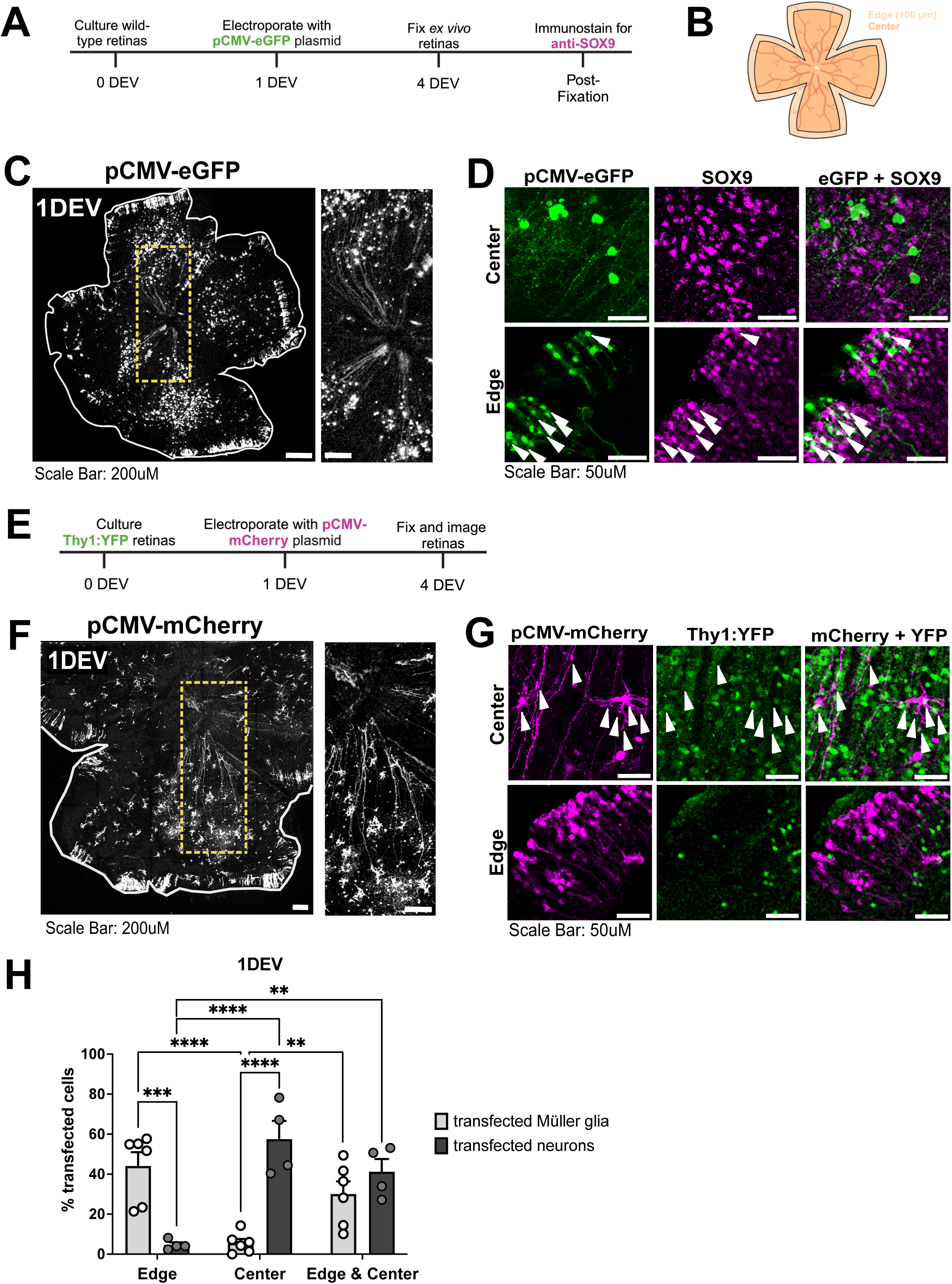
Preferential transfection at 1DEV of Müller glia and neurons at the edge and center of tissue, respectively. (A) Timeline of experimental design for 1DEV transfection of wild type tissue with pCMV-eGFP reporter plasmid and staining for SOX9 to detect Müller glia. (B) Schematic indicating edge (100µm from the retina periphery) and center of tissue. (C) pCMV-eGFP transfected cells in wild-type tissue transfected at 1DEV. Scale bar: Left, 200µm. Right, 50µm. (D) Left: pCMV-eGFP transfected cells, middle: SOX9 staining, and right: overlap of eGFP and SOX9 at the center (top) and edge (bottom) of tissue at 1DEV. Arrows indicate double-labeled cells. Scale Bar: 50µm.(E) Timeline of experimental design for 1DEV transfection of Thy1:YFP neuronal reporter tissue with pCMV-mCherry reporter plasmid. (F) pCMV-mCherry transfected cells in Thy1:YFP tissue transfected at 1DEV (YFP not shown). Scale bar: Left, 200µm, Right, 50µm. (G) Left: pCMV-mCherry transfected cells, middle: Thy1:YFP reporter, and right: overlap of mCherry and YFP at the center (top) and edge (bottom) of tissue after electroporation at 1DEV. Arrows indicate double-labeled cells. Scale Bar: 50µm. (H) Quantification of percentage of cells transfected with pCMV-eGFP that are SOX9+ and cells transfected with pCMV-mCherry that are Thy1:YFP+ at the edge, center and combined edge and center of tissue after electroporation at 1DEV. Mean ± SEM is shown. N=4-6. *p < 0.05, **p < 0.01, ***p < 0.001, ****p < 0.0001. No bracket indicates p > 0.05. All statistical tests performed on arcsine transformed values.

### Müller glia are preferentially transfected at 14DEV

Changes in cell survival, location, size, and morphology occur over time in culture^10–12^. Retinal tissue was electroporated at 14DEV to assess differences in transfection profile over time. Cells transfected at the edge of tissue resembled Müller glial morphology (Figure 3D, G), but the characteristic retinal ganglion cell morphology visible in the center in retinas transfected at 1DEV was not observed (Figure 3C, D, F, G). Staining for SOX9 revealed an increase in transfected Müller glia compared to inner retinal neurons as assessed with Thy1:YFP regardless of location (Figure 3D, G). These observations suggest that there is a decrease in neurons transfected at 14DEV overall, and an increase in transfection of Müller glia at the tissue center (Figure 3H).

**Figure 3.**
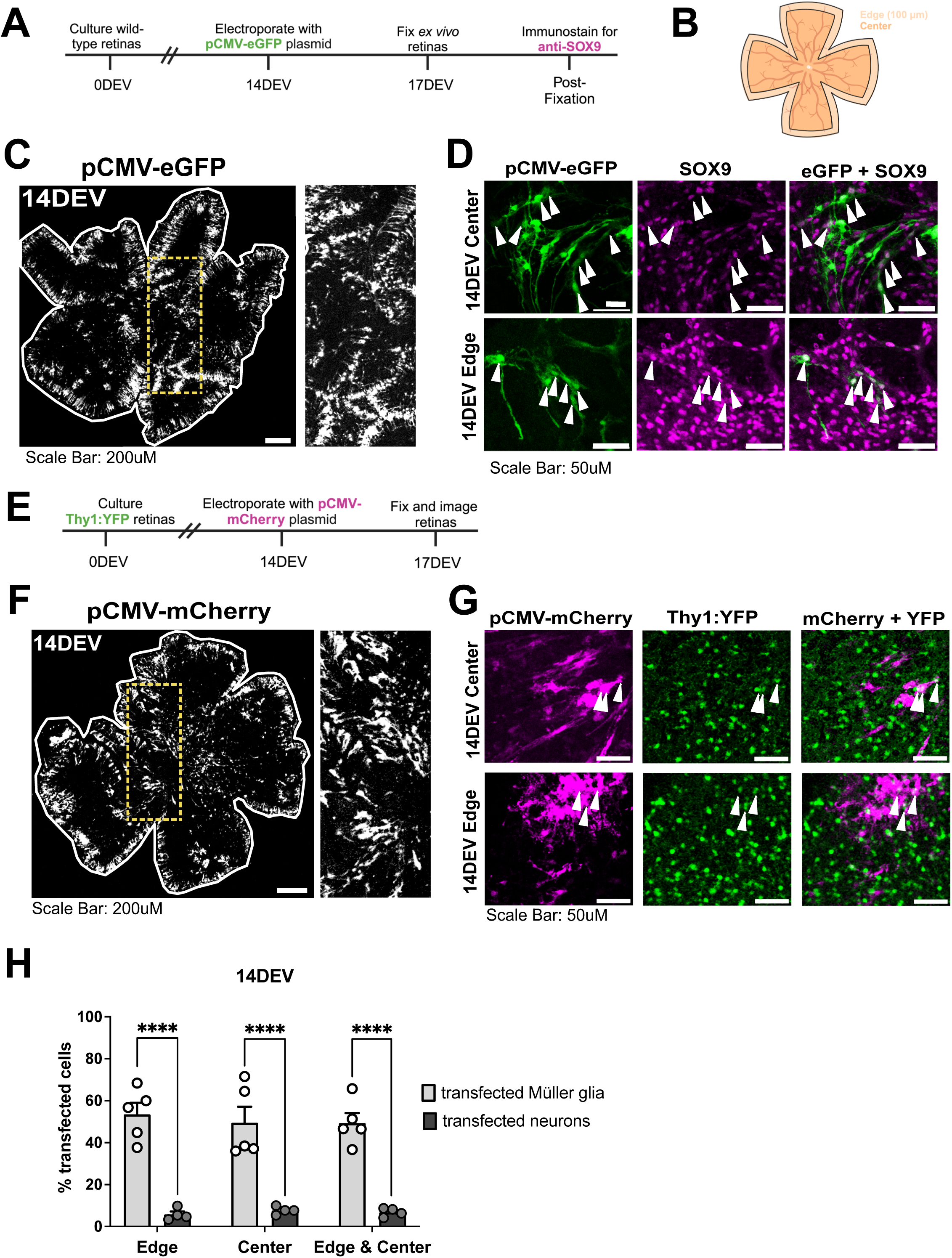
Preferential transfection of Müller glia at 14DEV. (A) Timeline of experimental design for 14DEV transfection of wild type tissue with pCMV-eGFP reporter plasmid and staining for SOX9 to detect Müller glia. (B) Schematic indicating edge (100µm from the retina periphery) and center of tissue. (C) pCMV-eGFP transfected cells in wild-type tissue transfected at 14DEV. Scale bar: Left, 200µm. Right, 50µm.(D) Left: pCMV-eGFP transfected cells, middle: SOX9 staining, and right: overlap of eGFP and SOX9 at the center (top) and edge (bottom) of tissue at 14DEV. Arrows indicate double-labeled cells. Scale Bar: 50µm. (E) Timeline of experimental design for 14DEV transfection of Thy1:YFP neuronal reporter tissue with pCMV-mCherry reporter plasmid. (F) pCMV-mCherry transfected cells in Thy1:YFP tissue transfected at 14DEV (YFP not shown). Scale bar: Left 200µm, Right, 50µm. (G) Left: pCMV-mCherry transfected cells, middle: Thy1:YFP reporter, and right: overlap of mCherry and YFP at the center (top) and edge (bottom) of tissue after electroporation at 14DEV. Arrows indicate double-labeled cells. Scale Bar: 50µm. (H) Quantification of percentage of cells transfected with pCMV-eGFP that are SOX9+ and cells transfected with pCMV-mCherry that are Thy1:YFP+ at the edge, center and combined edge and center of tissue after electroporation at 14DEV. Mean ± SEM is shown. N=4-5. *p < 0.05, **p < 0.01, ***p < 0.001, ****p < 0.0001. No bracket indicates p > 0.05. All statistical tests performed on arcsine transformed values.

To visualize tissue structure with respect to inner neurons and Müller glia at 1 and 14DEV, orthogonal slice projections were constructed from Z-stack confocal images of untransfected Thy1:YFP or Rlbp1:eGFP transgenic retinas, the latter serving as a reporter for Müller glia^32^. An ROI “slice” was cropped from each image in the same orientation to observe center and edge regions (Figure 4). At 1DEV, YFP+ cells were observed in the ganglion cell layer (GCL) and inner nuclear layer (INL; Figure 4A). Fluorescence was also observed in the inner plexiform layer (IPL), suggesting maintained laminar organization. However, fewer YFP+ cells were observed in the GCL at the retinal edge (Figure 4A, left). At 14DEV, YFP expression in the GCL was diminished in the central region but interestingly was retained in the IPL and INL (Figure 4B). In Rlbp1:eGFP tissue, GFP expression was also detectable in the GCL at the center of tissue at 1DEV (Figure 4C), consistent with their radial morphology^33^. At 14DEV, there was an increase in GFP expression relative to the total tissue volume including at the basal surface of the retina, where plasmids are placed for electroporation (Figure 4D). This could be due to Müller glia survival, proliferation, hypertrophy, altered morphology, or increased GFP expression on a per cell basis, none of which are mutually exclusive. Importantly, the loss of RGCs and accumulation of Müller glia at the basal surface could explain the shift toward preferential transfection of Müller glia at 14DEV.

**Figure 4.** Changes in tissue structure at electroporation surface in orthogonal slice projections of 1DEV & 14DEV Thy:YFP and Rlbp1:eGFP retinas favor electroporation of Müller glia at 14DEV. Orthogonal slice projections of center and edge created from 3D projections of ROIs (354.25 pixels by 11.76 pixels) to visualize cross sections within *ex vivo* retinas from transgenic mouse models. ONL: Outer nuclear layer. INL: Inner nuclear layer. GCL: Ganglion cell layer. IPL: Inner plexiform layer. * denotes the optic nerve head (ONH). (A) Orthogonal slice projections of the edge (left) and center (right) of Thy1:YFP *ex vivo* tissue fixed at 1DEV. Scale Bar:100µm. (B) Orthogonal slice projections of the edge (left) and center (right) of Thy1:YFP *ex vivo* tissue fixed at 14DEV. C) Orthogonal slice projections of the edge (left) and center (right) of Rlbp1:eGFP *ex vivo* tissue fixed at 1DEV. (D) Orthogonal slice projections of the edge (left) and center (right) of Rlbp1:eGFP *ex vivo* tissue fixed at 14DEV. (N=4).

### Electrical stimulation induces BrdU incorporation in *ex vivo* retina

The adult mouse retina is normally quiescent, but proliferation can occur to a limited extent in non-neuronal cells such as microglia and Müller glia in response to injury and/or inflammation, two contexts that are likely occurring in the *ex vivo* retina ^6,10,12,34–39^. Proliferation has also been shown to increase in response to electrical stimulation^40–44^, though this has yet to be shown in the retina. We therefore assessed proliferation in the cultures with and without electrical stimulation. Electrical stimulation was delivered using the electroporator with the same settings for transfection and three conditions were tested: no electroporation (-*EP*), electroporation with DNA loading solution lacking DNA (+*EP*), or electroporation with pCMV-eGFP plasmid (*+EP^DNA^*). These treatments were applied to *ex vivo* retinas at 1DEV or 14DEV. At 72 hours post-treatment,16uM BrdU was delivered in the cell culture media for 24 hours to assess cell cycle reentry that persists beyond the initial response to electrical stimulation (Figure 5A). At both times in culture, there were significant increases in BrdU+ cells per mm^3^ in the +*EP* and *+EP^DNA^* conditions compared to the -*EP* condition (Figure 5B,C). Similar levels of BrdU incorporation in the +*EP* and *+EP^DNA^*conditions were anticipated since eGFP expression is not expected to promote proliferation. The increase in proliferation observed in +*EP* and *+EP^DNA^* conditions compared to the -*EP* condition revealed that electrical stimulation of retinal tissue *ex vivo* can lead to changes in cell behavior associated with cell cycle activity.

**Figure 5.**
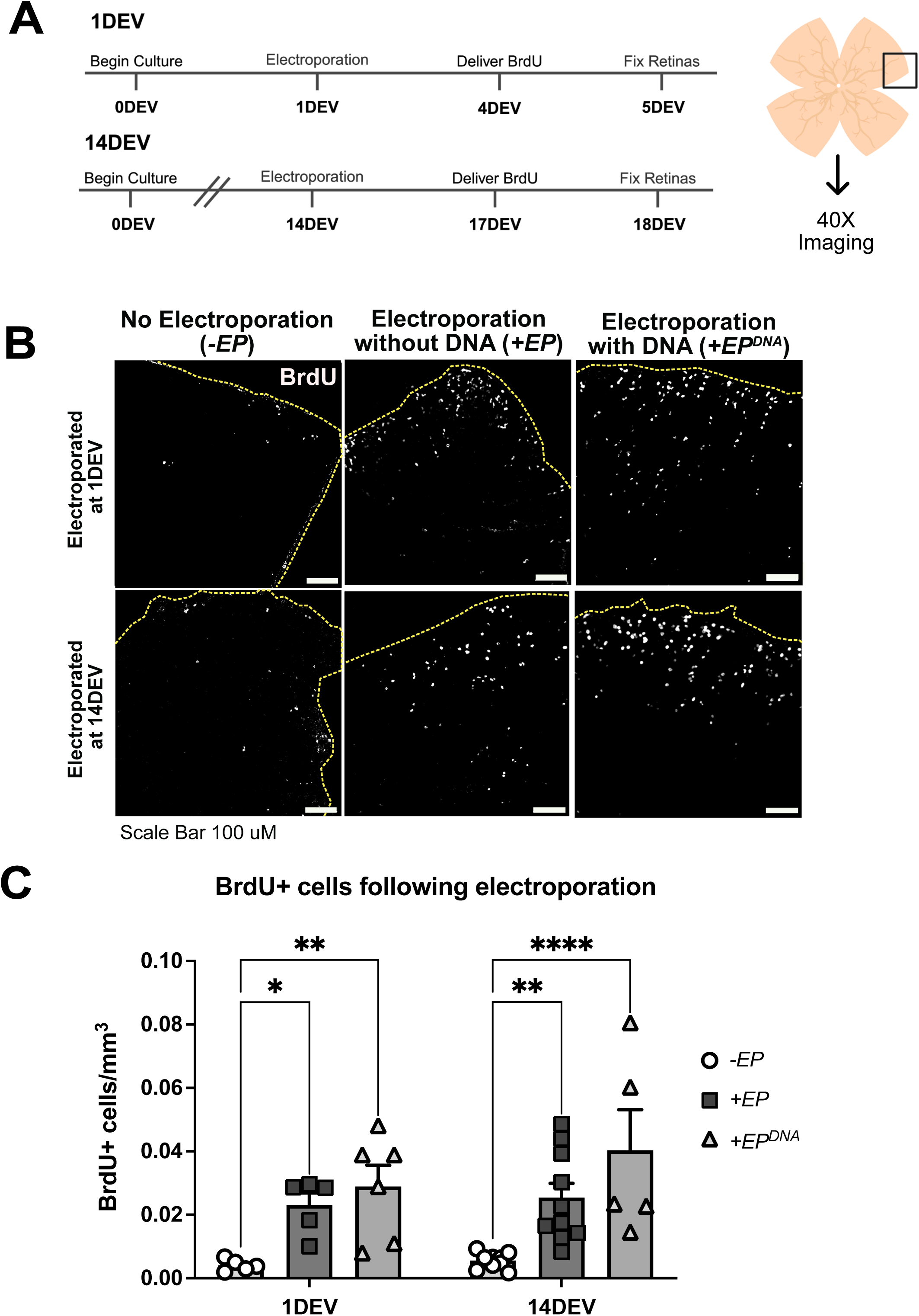
BrdU incorporation increases with electrical stimulation in the *ex vivo* retina. (A) Timeline of experimental design for tissue electroporated at 1DEV and 14DEV. BrdU is delivered in culture medium for 24 hours. Following fixation, pieces of retinal tissue were cut, stained for BrdU incorporation, and imaged using confocal microscopy. No Electroporation (*-EP*) retinas were given BrdU and fixed at the same timepoints indicated for electroporated retinas (*+EP, +EP^DNA^*). (B) BrdU incorporation *- EP* (left), *+EP* (center) and *+EP^DNA^* (right) retinas at 1DEV (top) and 14DEV (bottom). Scale Bar: 100µm. (C) Quantification of the number of BrdU+ cells detected per tissue volume (mm^3^). Mean ± SEM is shown. N=5-10. *p < 0.05, **p < 0.01, ***p < 0.001, ****p < 0.0001. No bracket indicates p > 0.05. All statistical tests performed on arcsine transformed values.

### The proportion of BrdU+ cells that are Müller glia increases in 14DEV compared to 1DEV retinas and following electrical stimulus

We predicted that microglia and Müller glia were re-entering the cell cycle in response to electrical stimulation. To assess this, *ex vivo* retinas were immunostained with the microglial marker IBA1 or with SOX9 in combination with BrdU detection, and colocalization with BrdU was quantified to determine the number of these cells undergoing cell cycle entry (Figure 6). In retinas treated at 1DEV or 14DEV, BrdU+ microglia and Müller glia were observed in *-EP* and *+EP* conditions (Figure 6A, B). Quantification revealed a significantly increased proportion of BrdU+ Müller glia, indicated by SOX9 labeling, in retinas treated at 14DEV in *+EP* retinas compared to 1DEV *+EP* (Figure 6C). The number of BrdU+ Müller glia significantly increased in *+EP* retinas treated at 14DEV compared to 1DEV *+EP* retinas and 14DEV *-EP* retinas, whereas the number of BrdU+ microglia remained steady (Figure 6D). These results suggest an increase in cell cycle entry in Müller glia that correlates with time in culture and electrical stimulation.

**Figure 6.**
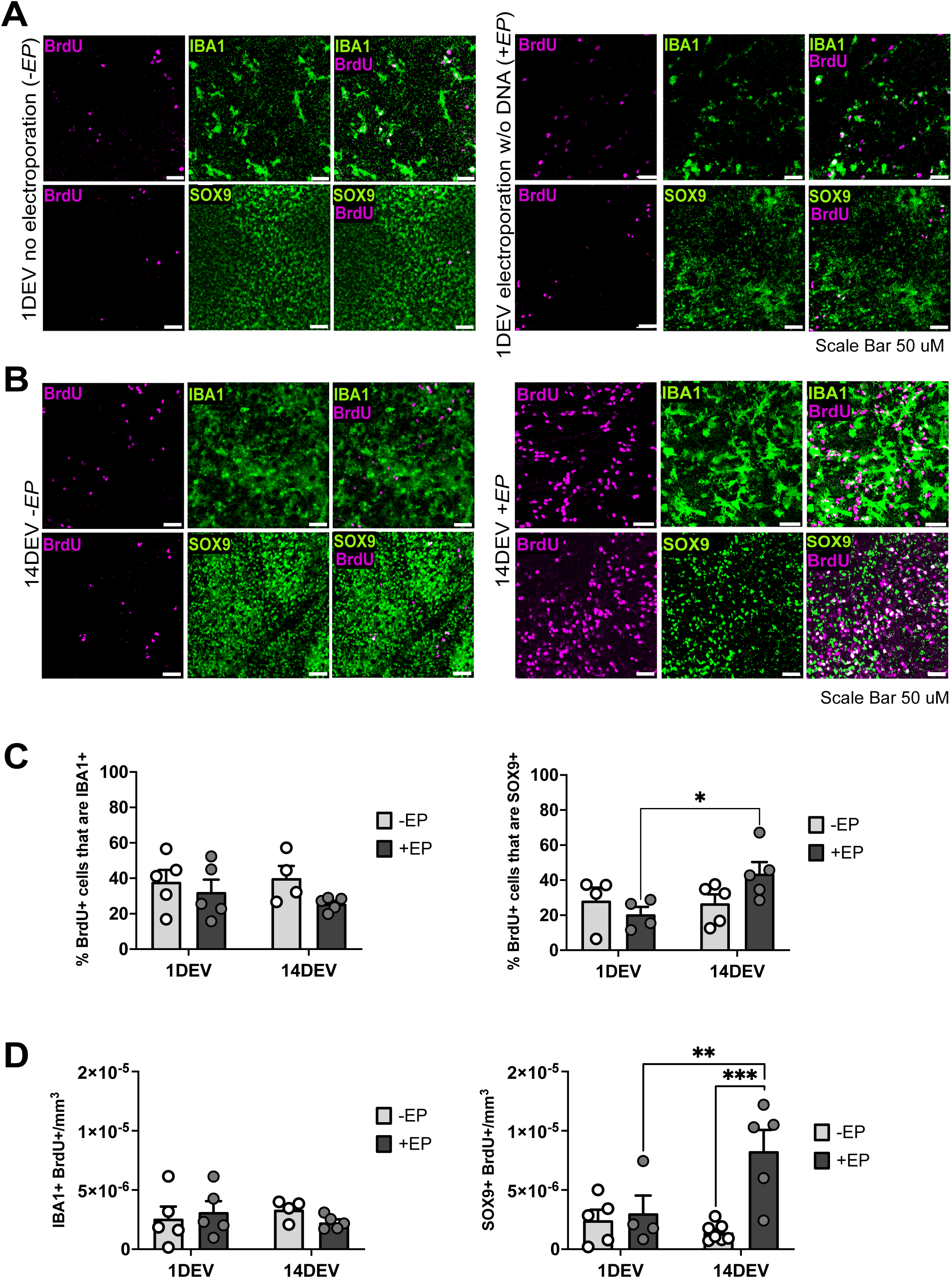
Increase in BrdU+ Müller glia at 14DEV vs. 1DEV and following electrical stimulus. (A) 1DEV *-EP* (left) or *+EP* (right) retinas stained for IBA1 (top left) or SOX9 (bottom left) and BrdU (top middle & bottom middle). Overlap of IBA1 & BrdU (top right) or SOX9 & BrdU (bottom right). Arrows indicate double labeled cells. Scale bar: 50µm. (B) 14DEV *-EP* (left) or *+EP* (right) retinas stained for IBA1 (top left) or SOX9 (bottom left) and BrdU (top middle & bottom middle). Overlap of IBA1 & BrdU (top right) or SOX9 & BrdU (bottom right). Colabeled cells appear white. Scale bar: 50µm. (C) Quantification of the percent of BrdU+ cells that are IBA1+ (left) or SOX9+ (right) in *-EP* and *+EP* conditions at 1DEV and 14DEV. D) Quantification of colabeled IBA1+ (left) or SOX9+ (right) cells colabeled as BrdU+ per tissue volume (mm^3^) in *-EP* and *+EP* conditions at 1DEV and 14DEV. Mean ± SEM is shown. N=5-10. *p < 0.05, **p < 0.01, ***p < 0.001, ****p < 0.0001. No bracket indicates p > 0.05. All statistical tests performed on arcsine transformed values.

### Transfection of pCMV-Cre reveals a higher transfection efficiency than pCMV-eGFP

A limitation of using fluorescent reporter expression constructs such as pCMV-eGFP is that cells could have been transfected but expression of the fluorescent protein was below detection. Since as few as 4 molecules of Cre recombinase are sufficient for recombination^45–47^, we predicted that transfection of a construct expressing Cre under the control of the same promoter/enhancer elements (pCMV-Cre) would reveal a higher transfection efficiency than what can be detected with pCMV-eGFP (Figure 7A). To test this, pCMV-Cre^48^ and pCMV-eGFP were co-transfected at equimolar concentrations into *ex vivo* retinas from *Rosa^AI14^* mice, which express tdTom after Cre recombination*. Ex vivo* retinas were analyzed 5 days after transfection at 1DEV or 14DEV.

**Figure 7.**
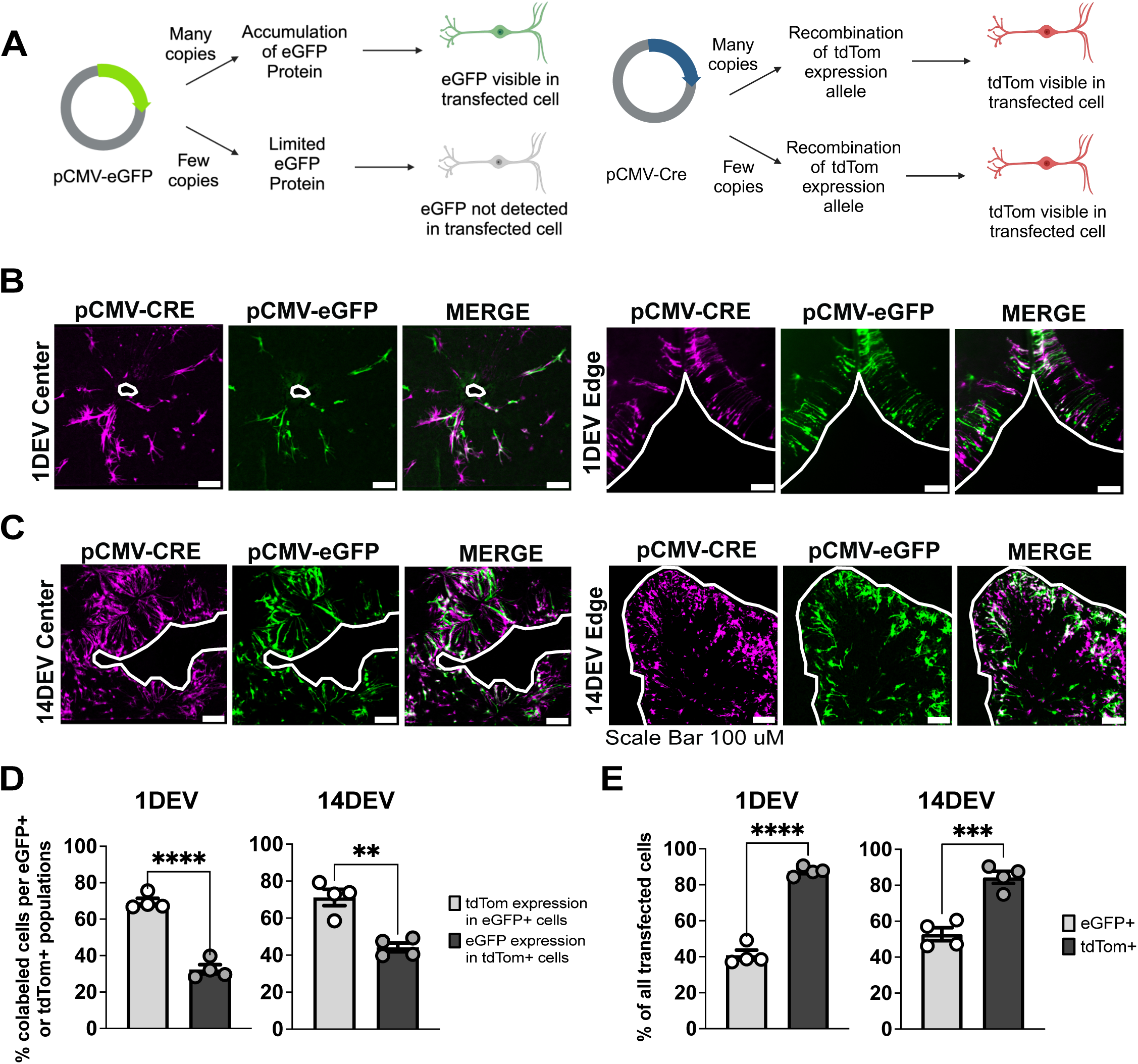
Co-transfection pCMV-eGFP and pCMV-Cre reveals increased detectability of transfected cells with a Cre expression construct. (A) Schematic of detection of pCMV-eGFP expression (left) vs. pCMV-Cre expression (right) in a *Rosa^AI14^* reporter tissue when few or many copies of the construct are transfected in a cell. (B) Cells transfected with equimolar concentration of pCMV-CRE and pCMV-eGFP at 1DEV at the center (left) or edge (right). Scale bar: 100µm. (C) Cells transfected with equimolar concentration of pCMV-CRE and pCMV-eGFP at 14DEV at the center (left) or edge (right) of retinas electroporated at 14DEV with both constructs at equimolar concentration. Scale bar: 100µm. (D) Quantification of the proportion (%) of colabeled cells detectable as transfected with pCMV-eGFP (eGFP+) and pCMV-Cre (tdTom+) within eGFP+ or tdTom+ populations transfected at 1DEV (left) and 14DEV (right).. (E) Quantification of the proportion (%) of cells detectable as transfected with pCMV-eGFP (eGFP+) and pCMV-Cre (tdTom+) within the population of all cells transfected at 1DEV (left) and 14DEV (right). Mean ± SEM is shown. N=4. *p < 0.05, **p < 0.01, ***p < 0.001, ****p < 0.0001. No bracket indicates p > 0.05. All statistical tests performed on arcsine transformed values.

As expected, we observed similar expression patterns of eGFP and tdTom in co-transfected retinas, regardless of location in the tissue or days in culture (Figure 7B, C). Cell quantification revealed that the majority of eGFP+ cells expressed tdTom, but this was not the case for tdTom+ cells, of which the minority were GFP+ (Figure 7D). Consistent with this, tdTom+ cells constituted a higher proportion of the transfected population than eGFP+ cells (Figure 7E; Table S1). While several factors such as plasmid size, nucleic acid composition, or secondary structures could affect transfection efficiency, we expected the expression characteristics of these constructs to be similar since they are similar in size and share the same promoter and enhancer elements. In this context, these data suggest that the transfection efficiency is greater than what can be observed by transfection of expression plasmids for fluorescent proteins.

### pCMV-Cre exhibited the highest transfection efficiency when compared to other Cre-expression constructs

The CMV promoter is considered a cell ubiquitous promoter. However, studies have shown that CMV promoters are not expressed well in post-mitotic neurons compared to other cell types^49,50^ which could reduce the apparent transfection efficiency of CMV-based expression vectors in *ex vivo* retinal cultures. To address this, we tested two constructs where Cre is driven by distinct regulatory elements: pCAG-Cre which contains the promoter for the chicken *beta-Actin* gene and CMV immediate-early enhancer^22^, and pBS513 EF1alpha-Cre (pEf1α-Cre), which contains the promoter region of the *Elongation Factor 1 alpha* gene^51^. In each case, pCMV-eGFP was co-transfected at equimolar concentrations. pCAG-Cre and pEf1α-Cre activated tdTom expression and the expression patterns of tdTom overlapped with eGFP (Figure 8A, D). Cell quantification of retinas co-transfected at 1DEV with pCAG-Cre and pCMV-eGFP revealed similar proportions of eGFP+ cells expressing tdTom and tdTom+ cells expressing eGFP (Figure 8B, left), whereas in retinas transfected at 14DEV, the proportion of eGFP+ cells expressing tdTom was greater than tdTom+ cells expressing eGFP (Figure 8B, right). At both timepoints, tdTom+ cells constituted a higher proportion of the entire transfected cell population (Figure 8C). These data suggest pCAG-Cre behaves similarly to pCMV-Cre.

**Figure 8.**
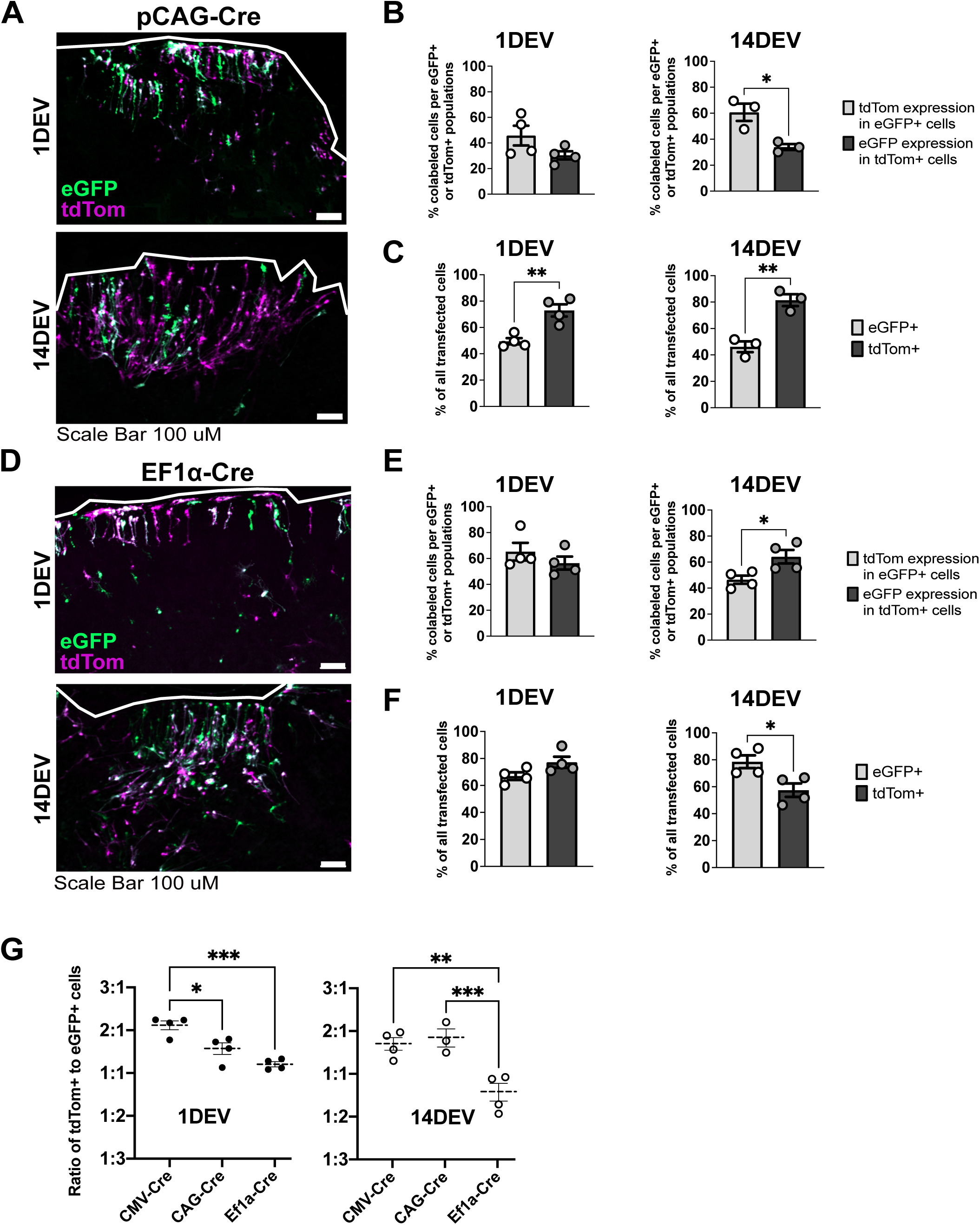
Variation in transfection efficiency in constructs with different expression elements. (A) Cells transfected with equimolar concentration of pCAG-Cre and pCMV-eGFP at 1DEV (top) and 14DEV (bottom) Scale bar: 100µm. (B) Quantification of the proportion (%) of colabeled cells detectable as co-transfected with pCMV-eGFP (eGFP+) and pCAG-Cre (tdTom+) within total eGFP+ or tdTom+ populations at 1DEV (left) and 14DEV (right). C) Quantification of the proportion (%) of cells detectable as transfected with pCMV-eGFP (eGFP+) and pCAG-Cre (tdTom+) within the population of all transfected cells at 1DEV (left) and 14DEV (right). (D) Cells transfected with equimolar concentration of pEF1α-Cre and pCMV-eGFP at 1DEV (top) and 14DEV (bottom) Scale bar: 100µm. (E) Quantification of the proportion (%) of colabeled cells detectable as co-transfected with pCMV-eGFP (eGFP+) and pEF1α-Cre (tdTom+) within total eGFP+ or tdTom+ populations at 1DEV (left) and 14DEV (right). (F) Quantification of the proportion (%) of cells detectable as transfected with pCMV-eGFP (eGFP+) and pEF1α-Cre (tdTom+) within the population of all transfected cells at 1DEV (left) and 14DEV (right). (G) Quantification of ratio of tdTom+ cells to eGFP+ cells at 1DEV and 14DEV in retinas co-transfected with pCMV-eGFP and either pCMV-Cre, pCAG-Cre or pEF1α-Cre. Mean ± SEM is shown. N=3-4. *p < 0.05, **p < 0.01, ***p < 0.001, ****p < 0.0001. No bracket indicates p > 0.05. Statistical tests performed on arcsine transformed values in B, C, E and F. The Y-Axis is Log_2_ transformed in G.

Like pCAG-Cre, retinas co-transfected at 1DEV with pEF1α-Cre and pCMV-eGFP had similar proportions of eGFP+ cells expressing tdTom and tdTom+ cells expressing eGFP (Figure 8E, left). In contrast to pCAG-Cre, the proportion of tdTom+ cells expressing eGFP was greater than eGFP+ cells expressing tdTom in retinas transfected at 14DEV (Figure 8E, right). These patterns were also reflected in the relative proportions of eGFP+ or tdTom+ cells that comprise the transfected cell population (Figure 8F).

Next, we compared the transfection efficiencies of pCMV-Cre, pCAG-Cre, and pEF1α-Cre. Since pCMV-eGFP was co-transfected with each construct in equimolar quantities, we calculated the ratios of tdTom+ cells to eGFP+ cells in the co-transfected retinas to compare the relative transfection efficiencies of the three Cre expression constructs. In retinas transfected at 1DEV, pCMV-Cre was most efficient at activating tdTom expression, as revealed by a ratio of 2.4 tdTom+ cells per eGFP+ cell and compared to pCAG-Cre at 1.5 tdTom+ cells per eGFP+ cell, and pEF1α-Cre at 1.1 tdTom+ cells per eGFP+ cell (Figure 8G, left; Table S1). At 14DEV, the efficiencies pCMV-Cre and pCAG-Cre were similar as revealed by ratios of 1.6 and 1.7 tdTom+ cells per eGFP+ cell, respectively. pEF1α-Cre was the least efficient at activating tdTom expression as indicated by a ratio of 1.1 eGFP+ cells per tdTom+ cell (Figure 8G, right). These results indicate that all three expression constructs can drive Cre expression in the *ex vivo* retina, but CMV regulatory elements are more efficient than the EF1α promoter. A comparison of the distributions of transfected cells expressing eGFP, tdTom, or both, for each construct supports this conclusion (Figure S1; Table S1).

## Discussion

This study introduces a novel method of electroporation for *ex vivo* retinal tissue cultured at the air-liquid interface that avoids submergence of the tissue or direct contact with the electrode. This electroporation method offers a rapid and time-dependent transfection efficiency while maintaining tissue integrity. This method is cost effective and does not require specialized equipment, making it generally accessible. Though we have only tested retinal tissue, it could easily be adapted for any tissue type that is cultured using an air liquid interface method, including lung, intestine, or skin^13,14,52,53^.

We observed a difference in the cell types transfected at different times in culture. This finding suggests that the timing of the transfection is useful for targeting specific cell populations within the retina. Müller glia undergo hypertrophy and migration in response to injury signals from retinal neurons^54–58^. Likely, the increase in availability of Müller glia at the electroporation surface is due to the loss of retinal ganglion cells and changes in Müller glial morphology associated with tissue remodeling^59^.

We found that the application of electrical stimulus alone leads to an increase in proliferation in Müller glia as indicated by BrdU incorporation. The mammalian retina is quiescent after retinal development, but Müller glia and microglia are known to exhibit limited proliferation in response to injury or inflammation. This proliferation occurs typically in young postmitotic mice and diminishes after injury onset^6,10,12,34–36,39,55,58^. Postmitotic Müller glia have been shown to proliferate in culture, however this effect was observed in *ex vivo* retinas from P10 mice and diminished by P14^10^. The mice used here were adults between 8-12 weeks of age.

Proliferation resulting from electrical stimulation has been reported in multiple cell types, including neural stem cells, human embryonic stem cells, osteoblasts, and fibroblasts^40,64–67^. Electroporation can also stimulate neural progenitor cell differentiation into neurons^67–69^. Amongst the BrdU+ cell population, there was a significant increase in BrdU+ Müller glia at 14DEV and following electrical stimulus whereas the presence of BrdU+ microglia was not dependent on time in culture. This could indicate an increase in Müller glia in the tissue at 14DEV or a change in cell state that leads to increased proliferation of Müller glia. Further studies are needed to determine whether proliferating Müller glia in this context also acquire neurogenic properties.

Delivery of a Cre expression construct into a recombination reporter mouse strain predicts that true transfection efficiency is greater than observed with a fluorescent reporter expression construct, meaning more cells may receive gene delivery than accounted for. It is advantageous with this system to use a sensitive reporter for electroporation, like Cre-mediated reporter expression, or to boost the signal for the fluorescent reporter with antibodies to enhance detection when analyzing fixed tissue.

Clinical studies indicate that low current electrical stimulation to the eye leads to better outcomes in patients with retinal dystrophies^60,61^. *In vitro* studies in Müller glia isolated from the dissociated mouse retina show that electrical stimulation increases expression of neurotropic factors and retinal progenitor markers, leading to an increase in proliferation^62,63^. The effect of electrical stimulation on Müller glia in the intact mouse retina has not been shown. This indicates a change in cell behavior following electrical stimulus that is not typical under normal physiological conditions.

## Limitations of Study

While this study provides valuable insights into the overall efficiency of transfection, especially on some retinal cell types, further research is necessary to fully characterize the effects on every cell type. This method has only been tested on retinal tissue, though it could be adapted for other tissue types cultured in an air-liquid interface format. Proliferation observed in this study was measured using BrdU incorporation, which labeled cells during S-phase of mitosis. BrdU could incorporate in cells that are then arrested in S-phase and do not fully divide. Further testing is required to understand the extent of proliferation following electrical stimulation.

Although certain cell types may be more amenable to electroporation, the method is not inherently cell type specific. Other methods of transfection should be employed if a study requires expression to only occur in certain cell types. We demonstrated that BrdU incorporation is induced following electrical stimulation with this method. Based on this finding, it is important that studies that use electroporation to the *ex vivo* retina as a gene delivery method include electrical stimulation without DNA in addition to other transfection controls to distinguish the effects of electrical stimulation and genetic manipulation when assessing proliferation.

Comparisons of pCAG-Cre and pEF1α-Cre in this study do not account for variability in cell type-specific preferential transfection. It is outside the scope of this study to delineate cell type preferential transfection with these constructs but is worthy of investigation by those interested in using this method to target specific retinal cell types.

## Acknowledgments

Funding for this study was provided by the National Eye Institute (R01-EY013760; P30-EY008126), the Janet and Jim Ayers Research Fund in Regenerative Visual Science, the William A. Black Chair in Ophthalmology, and an unrestricted grant from Research to Prevent Blindness, Inc. M.L.S was supported by the Vanderbilt Vision Training Grant (T32-EY007135) and a Ruth L. Kirschstein Predoctoral Individual National Research Service Award (F31EY035554) from the National Eye Institute. Confocal imaging and Imaris software analysis were performed at the Vanderbilt Cell Imaging and Shared Resource. The Zeiss LSM710 confocal microscope used for this study was acquired with funding from NIH (S10-RR027396). We thank Zachary Sanchez for assistance in photographing the electroporation set up and to members of the Levine and Fuhrmann laboratories for their insights and feedback.

## Author Contributions

Conceptualization, E.M.L and M.L.S; Methodology E.M.L, H.H.L, and M.L.S; Formal Analysis, M.L.S; Investigation, M.L.S; Resources, E.M.L; Data Curation, M.L.S; Writing (original draft), M.L.S; Writing (review & editing), M.L.S, H.L., E.M.L; Visualization, M.L.S; Supervision, E.M.L; Project Administration, E.M.L; Funding Acquisition, E.M.L.

## STAR Methods

### I. Resource Availability

#### Lead Contact

Further information and requests for resources and reagents should be directed to EML.

#### Materials Availability

This study did not generate new unique reagents.

#### Data and code availability

Microscopy and quantification data reported in this paper will be shared upon request.

### II. Experimental model and participant details

#### Mice

B6/129 mice (Jackson Laboratory (#101043)): 8 weeks old, female.

Thy-1 YFP-16 (B6.Cg-Tg(Thy1-YFP)16Jrs/J) (Jackson Laboratory (#003709)): 8 to 12 weeks old, female and male.

Rlbp1-eGFP (Tg(Rlbp1-eGFP)1Eml (MGI: 6195229)): 8 to 12 weeks old, female and male.

Rlbp1-CreERT2; Rosa^Ai14^ (129S6.Cg-Tg(Rlbp1-cre/ERT2)1Tfur (MGI:Pending); B6.Cg- *Gt(ROSA)26Sor^tm14(CAG-tdTomato)Hze^*/J (Jackson Laboratory (#007914))), 8 to 12 weeks old, female and male.

All mice are co-housed in a 12h/12h light-dark cycle with water and food ad libitum. All transgenic mice are backcrossed to B/129 mice. All animal experiments with mice were approved under the protocol M1600235 by the Vanderbilt Institutional Animal Care and Use committee and conform to the ARVO guidelines for the use of animals in vision research. All possible efforts were made to minimize animal suffering and the number of animals used.

### III. Method Details

#### Tamoxifen administration

To activate Cre recombination in transgenic mice, 2 doses of 200µg per gram body weight (GBW) of tamoxifen (Sigma Aldrich, T2859) were delivered in corn oil (Sigma Aldrich, C8267) via oral gavage. Mice were euthanized for tissue collection at least 5 days after tamoxifen treatment.

#### Retinal dissection

Prior to tissue harvest, dissection tools were sterilized with 70% ethanol and UV light for 15 minutes, and workstation was sterilized with 70% ethanol. 1X HBSS with Ca^2+^ and Mg^2+^ (Gibco, 14025) with 10mM HEPEs (Sigma Aldrich, H0887) and 1X Antibiotics-Antimycotics (Gibco, 15240062) (referred to as HBSS hereon) was prepared and kept on ice. Mice were euthanized by CO_2_ asphyxiation and cervical dislocation. Each eye was enucleated, rinsed in cold HBSS, then transferred to a fresh HBSS dish. Under a dissection scope and on a cold plate to keep retinas chilled, an incision was made in the cornea. The cornea was removed using spring scissors. The sclera, choroid and RPE were removed, the optic nerve head was cut on the proximal side with spring scissors, leaving it intact in the retina, and the retina and lens were transferred to a fresh HBSS dish. The lens was removed. Using spring scissors, 4 incisions were cut into retinal tissue from periphery through 25-40% of tissue toward optic nerve head, creating a flower-shaped retinal tissue that can be flattened.

#### *Ex vivo* retinal explant tissue culture

On the day of culture, plating media with supplements was prepared: NeuroCult™ Plating Media (Stem Cell Technologies, 05713), 1X SM1 neuronal supplement (Stem Cell Technologies, 05711), 500µM L-Glutamine (Gibco, 25030149), and 100unit Penicillin/100µg Streptomycin (Pen/Strep) (ThermoFisher, 15140) (referred to as plating media hereon).

Retinal tissue was dissected as described above. A hydrophilic PTFE cell culture insert (Millicell, PICM0RG50) was prepared by pipetting 150uL of plating media onto insert membrane. Retinal tissue was mounted onto the PTFE membrane with the retinal apical surface (photoreceptor side) in juxtaposition. Tissue was placed into a 6-well plate (Corning, 353004) containing 1mL plating media and incubated at 37°C with 5% CO^2^. After 48 hours, 500uL of initial plating media was replaced with BrainPhys™ Neuronal Medium (Stem Cell Technologies, 05790) containing 1X SM1 neuronal supplement, 1X N2 Supplement-A (Stem Cell Technologies, 07152) and Pen/Strep (referred to as BrainPhys media hereon). 500µL of media was removed and replaced with fresh BrainPhys media every 48 hours for the extent of the culture.

#### Agarose Disk Electroporation

0.5% agarose (RPI, A20090) in 1X PBS was heated to dissolve. 500µL was pipetted into multiple wells of a 24-well plate (Corning, CLS3516) and left to solidify at room temperature for 30 minutes in a cell culture hood. An anode dish was made by lining a 100mm dish (Sigma Aldrich, Z358762) with aluminum foil, sterilized under UV light for 15 minutes, and connected to the electroporator (BTX, 45-0052) by clamping the anode cable onto the aluminum foil. Sterile, chilled PBS was pipetted onto the anode dish. Tools and workspace were sterilized with 70% ethanol. The PTFE insert containing the tissue culture was washed by gently dipping the membrane in fresh, sterile PBS. The PTFE insert was then placed on the PBS in the anode dish.

An arm electrode (Bulldog Bio, CUY700P7L) connected to the electroporator was mounted on a micromanipulator (Narishige, UMM-3C) with the electrode surface facing up. Using a plastic transfer pipet (Fisherbrand, 13-711-7M) cut above the tapered tip, an agarose disk was stamped from the solidified agarose and transferred using a spatula to the arm electrode face. The disk was gently pressed to ensure adherence without bubbles underneath.

The 12µL DNA-glycerol transfection solution was prepared using a combination of 3pMol plasmid DNA diluted in sterile PBS with 7.5% glycerol (Sigma Aldrich, C8267), and 2.5% methyl green dye (MCE, HYD0163). The DNA-Glycerol solution was pipetted on top of the agarose disk and the arm electrode was rotated 180° while the DNA-glycerol solution and the agarose disk stayed adhered via surface tension. The arm electrode was lowered until the DNA solution was in contact with the tissue. Once there is contact between all components of the system, a closed circuit is formed.

Each tissue was electroporated with five square wave pulses (25V) for 50ms each with 250ms intervals between pulses. The arm electrode was slowly raised to disconnect from the tissue, and the tissue was returned to the incubator. This protocol has only been tested with square wave pulses.

Expression construct pCMV-eGFP was generated by the Levine lab. pCMV-mCherry was a gift from Roger Tsien (ClonTech plasmid # 632524, https://www.takarabio.com/documents/Vector%20Documents/PT3975-5_060412.pdf). pCMV-Cre was a gift from David Liu (Addgene plasmid # 123133; http://n2t.net/addgene:123133; RRID:Addgene_123133). pCAG-Cre was a gift from Anjen Chenn (Addgene plasmid # 26647; http://n2t.net/addgene:26647; RRID:Addgene_26647), which was generated by Anjen Chenn by insertion of the Cre expression motif and deletion of the GFP expression motif into the backbone of pCAG-GFP, a gift from Connie Cepko (Addgene plasmid # 11150; http://n2t.net/addgene:11150; RRID:Addgene_11150). pBS513 EF1alpha-cre was a gift from Brian Sauer (Addgene plasmid # 11918; http://n2t.net/addgene:11918; RRID:Addgene_11918).

#### Retinal explant culture tissue fixation

Culture medium was removed and replaced with room temperature PBS to wash 3 times for 10 minutes each. 100µL 1X PBS was pipetted on top of tissue for each wash. Retinas were fixed in 4% PFA for 2 hours at room temperature or at 4**°**C for 24 hours. PFA was removed and fixed retinas were washed 3 times with 1 mL 1X PBS for 30 minutes at room temperature. Fixed retinas were stored at 4°C in 1 mL 1X PBS with 0.01% sodium azide (Sigma Aldrich, 08591).

#### Whole mount immunohistochemistry

Each quadrant of the retinal tissue was separated and cut in half to generate 8 tissue fragments for staining. Individual tissue fragments were transferred into a 96-well plate (Corning, CLS3628). Retinas were washed 3 times with 1mL 1X PBS containing 1% Triton-X (1% PBST), followed by 100µL 10% normal donkey serum (NDS) (SouthernBiotech, 003001) in 1% PBST (blocking buffer) for 2 hours at room temperature. Blocking buffer was removed and retinas were stained with primary antibodies in 1% NDS blocking buffer for 2 hours at room temperature or 24 hours at 4°C. Retinas were washed 3 times with 200µL 1% PBST for 30 minutes each. Secondary antibodies in 1% NDS blocking buffer were applied for 2 hours at room temperature. The secondary antibody was removed, and tissue was washed 3 times with 200µL 1% PBST for 30 minutes per wash. Stained retinas were mounted on Colorfrost plus slides (Fisherbrand, 12-550-17) in Fluoromount G (Invitrogen, 00-4958-02), coversliped, allowed to dry at room temperature overnight, then stored at 4°C until imaged.

The following primary antibodies were used: α-BrdU (1:200, Abcam, ab6326), α-IBA1 (1:500, FUJIFILM, 019-19741), and α-SOX9 (1:500, Chemicon, ab5535). Secondary antibodies conjugated to Alexa 488 (Invitrogen, A21206), Alexa 568 (Invitrogen, A10042), and Alexa 647 (Intvitrogen, 21247) were each diluted at 1:500.

#### BrdU treatment and detection

16µM BrdU was added to culture media at varying timepoints in culture per experiment. BrdU media was replenished daily until fixation. *Ex vivo* retinas were treated with BrdU for 24-72 hours depending on the experiment. Following treatment, tissues were fixed with PFA as described above. A 1/8 piece of tissue was selected at random and cut using as scalpel from the retinal culture to stain. Retinas were immunostained as described above for all other immunomarkers, then fixed with PFA for 30 min to preserve antigens prior to BrdU immunostaining. 2N HCl (Fisher Scientific, SA541) was used for antigen retrieval for 45 minutes at room temperature, followed by a 20-minute neutralization with 0.1M sodium borate (pH 8.5) (Millipore Sigma, 1066690010).

#### Imaging

Cultures were imaged using a fluorescent stereoscope (Nikon SMZ1270i) every 48 hours post-culture establishment, and every 24 hours post-electroporation monitor for morphological, structural, or fluorescent protein expression changes within the tissue. The Zeiss LSM 710 confocal microscope was used to image tissue cultures after fixation alone or fixation and staining. Images were analyzed using FIJI version 2.9.0.

#### Orthogonal slice projections

To observe structural changes in *ex vivo* retinal tissue cultures, 40X Z-Stack images were obtained from the Zeiss LSM 710 confocal microscope with 1µm steps between Z slices. Using FIJI version 2.9.0, 354.25 by 11.76 pixel ROIs were selected at random but angled from tissue periphery toward optic nerve head. These ROIs were converted to 3D projections and rotated to view as a cross section.

### IV. Quantification and Statistical Analysis

All cell counting and tissue volume calculations were calculated from 20X Z-stack, tile confocal images using Imaris 10.1 software. No statistical methods were used to predetermine sample sizes. GraphPad Prism 10 was used for the following statistical tests: All proportion values were arcsine transformed prior to statistical analysis. Comparisons between more than two groups were made by two-way ANOVA with Tukey post-hoc test. Comparisons between two groups within the same tissue, as in figure 7, were made using paired two-tailed t tests. Differences were considered statistically significant at p < 0.05. Data are presented as mean ± SEM. Statistical significance is indicated with asterisks: *p < 0.05, **p < 0.01, ***p < 0.001, ****p < 0.0001. Figure graphics created using Biorender.

## Supplemental Information

**Document S1. Figure S1 and Table S1**

**Figure S1: Proportion of eGFP+ and tdTom+ cells in tissues transfected with pCMV-eGFP and Cre expression construct with differing elements.** Euler plots of the proportion (%) of eGFP+, tdTom+, and eGFP+ tdTom+ colabeled cells per the sum of an edge and center 6.02×10^7^ um^3^ ROI. Retinas transfected at 1DEV (top) and 14DEV (bottom) with pCMV-eGFP and pCMV-Cre (left), pCAG-Cre (middle) or pEF1α-Cre (right). Euler plots are scaled to represent proportion. N=3-4. Statistical information available in Table S1.

**Table S1: Quantification of cells transfected at 1DEV and 14DEV with pCMV-eGFP and pCMV-Cre, pCAG-Cre or pEF1α-Cre.** Cell counts calculated from sum of cells detected in a 6.02×107 um3 region of interest at center and edge of each retinal sample. ROIs were selected based on presence of transfected cells. The negative inverse is shown for tdTom:eGFP ratios in which there were more eGFP+ cells than tdTom+ cells.

