## Supplementary figures and images for "Agarose disk electroporation method for *ex vivo* retinal tissue cultured at the air-liquid interface reveals electrical stimulus-induced cell cycle reentry in retinal cells"

### Supplemental Figure 1

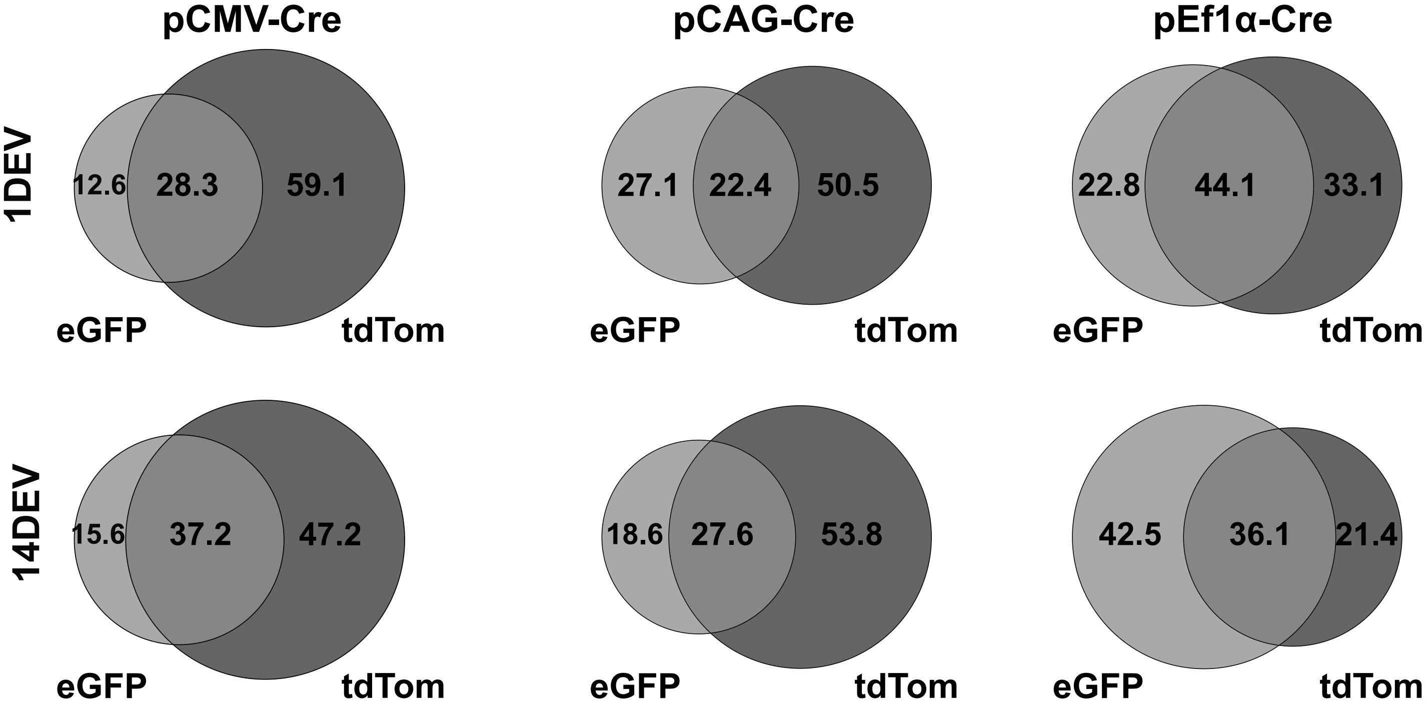
