## Supplemental Table 1 for "Agarose disk electroporation method for *ex vivo* retinal tissue cultured at the air-liquid interface reveals electrical stimulus-induced cell cycle reentry in retinal cells"

| Sample | Timepoint | Plasmids transfected | eGFP+ | tdTom+ | eGFP+ tdTom+ | Total eGFP+ | Total tdTom+ | Total transfected | eGFP+/Total transected (%) (Fig. S1) | tdTom+/Total transected (%) (Fig. S1) | eGFP+ tdTom+/Total transected (%) (Fig. S1) | tdTom+ in eGFP+ (%) (Figs. 7D,8B,8E) | eGFP+ in tdTom+ (%) (Figs. 7D,8B,8E) | Total eGFP/Total transected (%) (Figs. 7E,8C,8F) | Total tdTom/Total transected (%) (Figs. 7E,8C,8F) | Ratio tdTom:eGFP | Tranformed ratio tdTom:eGFP (Fig. 8G) |
|---|---|---|---|---|---|---|---|---|---|---|---|---|---|---|---|---|---|
| 1 | 1 DEV | pCMV-Cre + pCMV-eGFP | 33 | 216 | 102 | 135 | 318 | 351 | 9.4 | 61.5 | 29.1 | 75.6 | 32.1 | 38.5 | 90.6 | 2.4 | 2.4:1 |
| 2 | 1 DEV | pCMV-Cre + pCMV-eGFP | 29 | 138 | 58 | 87 | 196 | 225 | 12.9 | 61.3 | 25.8 | 66.7 | 29.6 | 38.7 | 87.1 | 2.3 | 2.3:1 |
| 3 | 1 DEV | pCMV-Cre + pCMV-eGFP | 31 | 100 | 66 | 97 | 166 | 197 | 15.7 | 50.8 | 33.5 | 68.0 | 39.8 | 49.2 | 84.3 | 1.7 | 1.7:1 |
| 4 | 1 DEV | pCMV-Cre + pCMV-eGFP | 26 | 134 | 53 | 79 | 187 | 213 | 12.2 | 62.9 | 24.9 | 67.1 | 28.3 | 37.1 | 87.8 | 2.4 | 2.4:1 |
| | | | | | | | | **MEAN** | ***12.6*** | ***59.1*** | ***28.3*** | ***69.3*** | ***32.4*** | ***40.9*** | ***87.4*** | ***2.2*** | ***2.2:1*** |
| | | | | | | | | **SEM** | ***1.3*** | ***2.8*** | ***2.0*** | ***2.1*** | ***2.6*** | ***2.8*** | ***1.3*** | ***0.2*** | ***0.2*** |
| **1** | 14DEV | pCMV-Cre + pCMV-eGFP | 127 | 543 | 358 | 485 | 901 | 1028 | 12.4 | 52.8 | 34.8 | 73.8 | 39.7 | 47.2 | 87.6 | 1.9 | 1.9:1 |
| 2 | 14DEV | pCMV-Cre + pCMV-eGFP | 65 | 102 | 92 | 157 | 194 | 259 | 25.1 | 39.4 | 35.5 | 58.6 | 47.4 | 60.6 | 74.9 | 1.2 | 1.2:1 |
| 3 | 14DEV | pCMV-Cre + pCMV-eGFP | 38 | 106 | 103 | 141 | 209 | 247 | 15.4 | 42.9 | 41.7 | 73.0 | 49.3 | 57.1 | 84.6 | 1.5 | 1.5:1 |
| 4 | 14DEV | pCMV-Cre + pCMV-eGFP | 89 | 500 | 340 | 429 | 840 | 929 | 9.6 | 53.8 | 36.6 | 79.3 | 40.5 | 46.2 | 90.4 | 2.0 | 2:1 |
| | | | | | | | | **MEAN** | ***15.6*** | ***47.2*** | ***37.2*** | ***71.2*** | ***44.2*** | ***52.8*** | ***84.4*** | ***1.6*** | ***1.6:1*** |
| | | | | | | | | **SEM** | ***3.4*** | ***3.6*** | ***1.6*** | ***4.4*** | ***2.4*** | ***3.6*** | ***3.4*** | ***0.2*** | ***0.2*** |
| 1 | 1 DEV | pCAG-Cre + pCMV-eGFP | 56 | 146 | 67 | 123 | 213 | 269 | 20.8 | 54.3 | 24.9 | 54.5 | 31.5 | 45.7 | 79.2 | 1.7 | 1.7:1 |
| 2 | 1 DEV | pCAG-Cre + pCMV-eGFP | 52 | 144 | 89 | 141 | 233 | 285 | 18.2 | 50.5 | 31.2 | 63.1 | 38.2 | 49.5 | 81.8 | 1.7 | 1.7:1 |
| 3 | 1 DEV | pCAG-Cre + pCMV-eGFP | 110 | 125 | 51 | 161 | 176 | 286 | 38.5 | 43.7 | 17.8 | 31.7 | 29.0 | 56.3 | 61.5 | 1.1 | 1.1:1 |
| 4 | 1 DEV | pCAG-Cre + pCMV-eGFP | 84 | 146 | 43 | 127 | 189 | 273 | 30.8 | 53.5 | 15.8 | 33.9 | 22.8 | 46.5 | 69.2 | 1.5 | 1.5:1 |
| | | | | | | | | **MEAN** | ***27.1*** | ***50.5*** | ***22.4*** | ***45.8*** | ***30.3*** | ***49.5*** | ***72.9*** | ***1.5*** | ***1.5:1*** |
| | | | | | | | | **SEM** | ***4.7*** | ***2.4*** | ***3.5*** | ***7.7*** | ***3.2*** | ***2.4*** | ***4.7*** | ***0.1*** | ***0.1*** |
| 1 | 14DEV | pCAG-Cre + pCMV-eGFP | 71 | 376 | 162 | 233 | 538 | 609 | 11.7 | 61.7 | 26.6 | 69.5 | 30.1 | 38.3 | 88.3 | 2.3 | 2.3:1 |
| 2 | 14DEV | pCAG-Cre + pCMV-eGFP | 103 | 184 | 94 | 197 | 278 | 381 | 27.0 | 48.3 | 24.7 | 47.7 | 33.8 | 51.7 | 73.0 | 1.4 | 1.4:1 |
| 3 | 14DEV | pCAG-Cre + pCMV-eGFP | 62 | 187 | 115 | 177 | 302 | 364 | 17.0 | 51.4 | 31.6 | 65.0 | 38.1 | 48.6 | 83.0 | 1.7 | 1.7:1 |
| | | | | | | | | **MEAN** | ***18.6*** | ***53.8*** | ***27.6*** | ***60.7*** | ***34.0*** | ***46.2*** | ***81.4*** | ***1.8*** | ***1.8:1*** |
| | | | | | | | | **SEM** | ***4.5*** | ***4.1*** | ***2.1*** | ***6.6*** | ***2.3*** | ***4.1*** | ***4.5*** | ***0.5*** | ***0.5*** |
| 1 | 1 DEV | pEF1α-Cre + pCMV-eGFP | 45 | 74 | 67 | 112 | 141 | 186 | 24.2 | 39.8 | 36.0 | 59.8 | 47.5 | 60.2 | 75.8 | 1.3 | 1.3:1 |
| 2 | 1 DEV | pEF1α-Cre + pCMV-eGFP | 15 | 37 | 88 | 103 | 125 | 140 | 10.7 | 26.4 | 62.9 | 85.4 | 70.4 | 73.6 | 89.3 | 1.2 | 1.2:1 |
| 3 | 1 DEV | pEF1α-Cre + pCMV-eGFP | 68 | 81 | 85 | 153 | 166 | 234 | 29.1 | 34.6 | 36.3 | 55.6 | 51.2 | 65.4 | 70.9 | 1.1 | 1.1:1 |
| 4 | 1 DEV | pEF1α-Cre + pCMV-eGFP | 66 | 76 | 100 | 166 | 176 | 242 | 27.3 | 31.4 | 41.3 | 60.2 | 56.8 | 68.6 | 72.7 | 1.1 | 1.1:1 |
| | | | | | | | | **MEAN** | ***22.8*** | ***33.1*** | ***44.1*** | ***65.3*** | ***56.5*** | ***66.9*** | ***77.2*** | ***1.2*** | ***1.2:1*** |
| | | | | | | | | **SEM** | ***4.2*** | ***2.8*** | ***6.4*** | ***6.8*** | ***5.0*** | ***2.8*** | ***4.2*** | ***0.0*** | ***0*** |
| 1 | 14DEV | pEF1α-Cre + pCMV-eGFP | 68 | 57 | 70 | 138 | 127 | 195 | 34.9 | 29.2 | 35.9 | 50.7 | 55.1 | 70.8 | 65.1 | -1.1 | 1:1.1 |
| 2 | 14DEV | pEF1α-Cre + pCMV-eGFP | 181 | 57 | 137 | 318 | 194 | 375 | 48.3 | 15.2 | 36.5 | 43.1 | 70.6 | 84.8 | 51.7 | -1.6 | 1:1.6 |
| 3 | 14DEV | pEF1α-Cre + pCMV-eGFP | 108 | 96 | 120 | 228 | 216 | 324 | 33.3 | 29.6 | 37.0 | 52.6 | 55.6 | 70.4 | 66.7 | -1.1 | 1:1.1 |
| 4 | 14DEV | pEF1α-Cre + pCMV-eGFP | 252 | 54 | 164 | 416 | 218 | 470 | 53.6 | 11.5 | 34.9 | 39.4 | 75.2 | 88.5 | 46.4 | -1.9 | 1:1.9 |
| | | | | | | | | **MEAN** | ***42.5*** | ***21.4*** | ***36.1*** | ***46.5*** | ***64.1*** | ***78.6*** | ***57.5*** | ***-1.4*** | ***1:1.4*** |
| | | | | | | | | **SEM** | ***5.0*** | ***4.7*** | ***0.5*** | ***3.1*** | ***5.2*** | ***4.7*** | ***5.0*** | ***0.2*** | ***0.2*** |
